# Genetic control of age-related gene expression and complex traits in the human brain

**DOI:** 10.1101/125195

**Authors:** Trevor Martin, Hunter B. Fraser

**Affiliations:** Department of Biology, Stanford University, Stanford, CA 94305, USA.

## Abstract

Age is the primary risk factor for many of the most common human diseases—particularly neurodegenerative diseases—yet we currently have a very limited understanding of how each individual’s genome affects the aging process. Here we introduce a method to map genetic variants associated with age-related gene expression patterns, which we call temporal expression quantitative trait loci (teQTL). We found that these loci are markedly enriched in the human brain and are associated with neurodegenerative diseases such as Alzheimer’s disease and Creutzfeldt-Jakob disease. Examining potential molecular mechanisms, we found that age-related changes in DNA methylation can explain some cis-acting teQTLs, and that trans-acting teQTLs can be mediated by microRNAs. Our results suggest that genetic variants modifying age-related patterns of gene expression, acting through both cis- and trans-acting molecular mechanisms, could play a role in the pathogenesis of diverse neurological diseases.

## Introduction

Despite the well-known importance of age in our understanding of diverse diseases, the molecular mechanisms by which aging exerts these effects are still mostly unknown (Niccoli and Partridge 2012). One key factor is likely to be gene expression regulation, the process by which the great diversity of cell types in every individual is generated from the same underlying genome. Indeed, many studies have reported widespread changes in gene expression during human aging, though the degree to which this dynamic regulation is affected by genetic differences between individuals is still mostly unknown. A well-studied exception is lactase persistence, which is a genetically-encoded age-associated phenotype. Genetic variants near the lactase gene can extend its expression from infancy to the entire human lifespan; these variants increase lactose digestion in an age-dependent manner, resulting in a selective advantage for adults who consume dairy products (Enattah et al. 2002).

More generally, age-related diseases could have a pathogenesis regulated at least in part by genomic loci that control gene expression over age. For example, the typical late in life onset of neurodegenerative diseases such as Alzheimer’s disease could potentially be related to the gradual up-or down-regulation of genes involved in its pathology over the human life span. Therefore individuals who have genotypes that protect against these regulatory changes could have decreased genetic risk for the disease.

Despite the known importance of age, most studies of the genetic control of gene expression across individuals (expression quantitative trait loci; eQTLs) either ignore age, or treat it as a confounding variable (GTEx Consortium 2015). Only a handful of studies have examined the role of genetic variation in human aging-related expression patterns (genotype-by-age interactions) (Kent et al. 2012; Yao et al. 2014; Glass et al. 2013; Brinkmeyer-Langford et al. 2016). These interactions may be especially important in the brain, since many neurodegenerative diseases have onsets that are typically late in life and whose risk could be affected by loci that control transcription over aging. The aging human brain is also interesting in light of theories of human evolution, such as the idea that neoteny, or developmental delay, may play a central role in human differentiation from close relatives such as chimpanzees (Somel et al. 2009). Indeed, age-related gene expression patterns differ markedly between human and chimpanzee cerebral cortex (Fraser et al. 2005).

In addition to mRNA expression levels, many other cellular phenotypes such as DNA methylation and microRNA expression could also be regulated by genotype-by-age interactions. Genetic control of DNA methylation over age has only been studied previously in human peripheral blood cells (van Dongen et al. 2016) and microRNA expression over age in dwarf mouse serum (Victoria et al. 2015). Here we report the first integrated analysis of age-dependent genetic regulation of mRNA expression, DNA methylation, and microRNA expression.

## Results

### Mapping loci affecting age-related gene expression

To identify genetic loci that control changes in expression over age, we calculated the significance of the interaction between SNP genotype and age on mRNA expression level in a genome-wide scan (see Methods). We call these loci temporal expression quantitative trait loci (teQTL; Fig. 1A; Supp. Fig. 1). We first applied this method to map teQTLs in 44 diverse tissues from hundreds of individuals (GTEx Consortium 2015). Surprisingly, we found that teQTLs were most prevalent in the brain (Wilcoxon p = 2.0x10^-3^; Fig. 1B), with other tissues having very few significant teQTLs, despite many of these having more sampled individuals (and thus more power to detect teQTLs). This teQTL brain enrichment stands in stark contrast to traditional eQTLs, for which the number of samples and the number of expressed genes are the major determinants of the number of detected eQTLs (Supp. Fig. 2). The brain regions with the highest enrichment for teQTLs are the caudate basal ganglia and frontal cortex.

**Figure 1.**
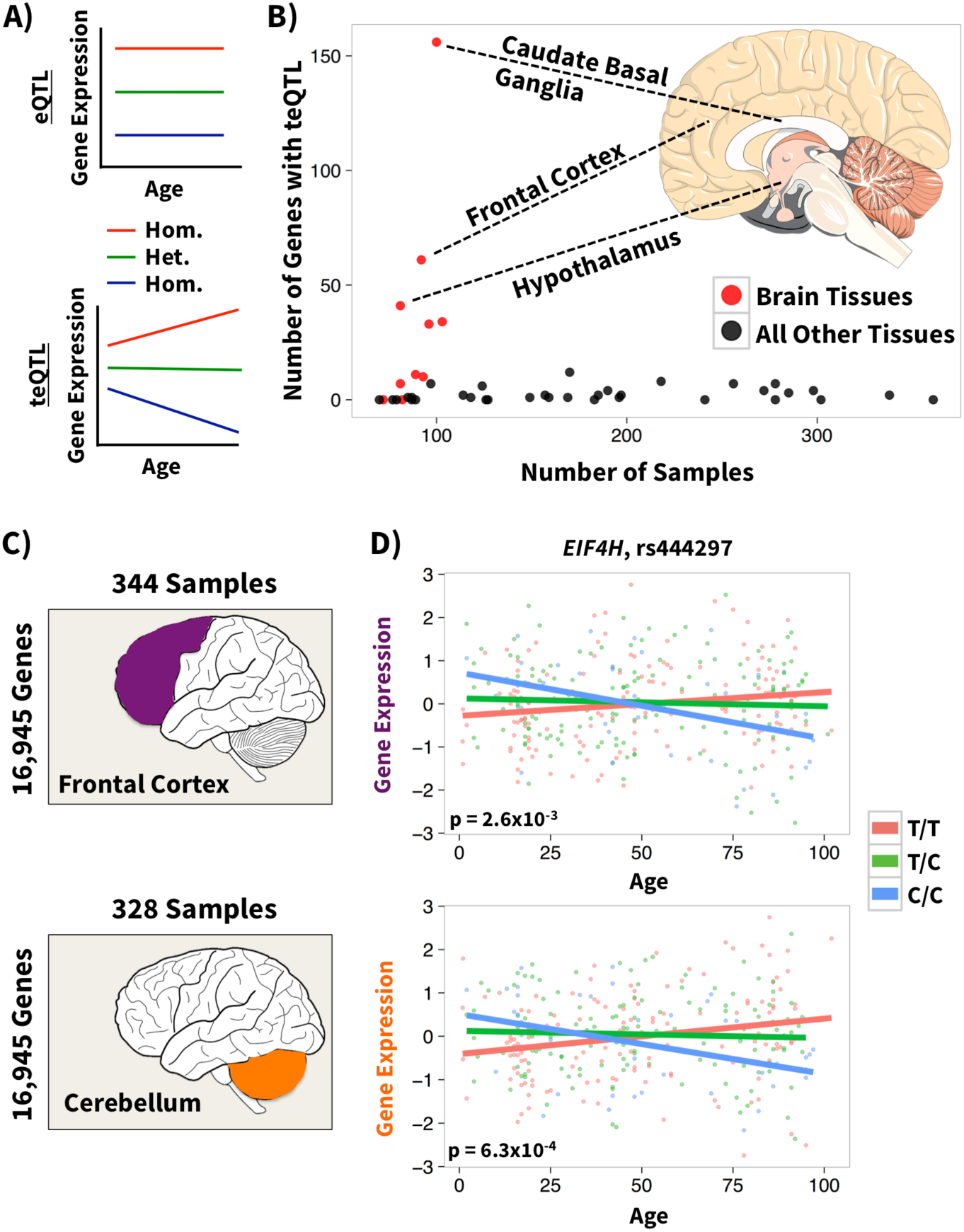
Identification of teQTLs. (A) Overview of the teQTL method. In traditional eQTL analysis differences in mean gene expression across individuals with each of the three possible genotypes at a nearby SNP are detected. In the teQTL analysis interactions between genotype and age that influence gene expression level are detected. (B) The number of genes whose expression has a significant age x genotype interaction plotted versus the number of individuals profiled for each tissue in the GTEx data set. In red are all tissues from the brain (10), in black are all other tissues (34). Dotted lines connect selected tissues to their location in the brain. (C) Overview of mRNA expression data. 344 individuals with frontal cortex data (purple) and 328 individuals with cerebellum (orange) gene expression data were used in this analysis. (D) Example of a teQTL present in both tissues. Normalized gene expression mRNA levels are shown on the y-axis for both the frontal cortex (top; *β* _age*geno_ = -0.01) and cerebellum (bottom; *β* _age*geno_ = -0.01) versus age for the gene *EIF4H*. A linear regression fit for each of the three genotypes at the local SNP rs444297 are shown in the colors red (major allele homozygote), green (heterozygote), and blue (minor allele homozygote).

In order to explore these brain teQTLs further, we applied our analysis to a larger independent data set of mRNA expression in 344 frontal cortex samples and 328 cerebellum samples, with ages ranging from 1 to 102 years old (mean age of 50 years; Fig. 1C; hereafter the “primary” data set) (Gibbs et al. 2010). At an FDR of 20% we identified 44 teQTLs in frontal cortex, and 85 in cerebellum (Supp. Tables 1-2; Supp. Figure 3). Overall, 53% of teQTLs were also nominally significant (p < 0.05) eQTLs, and 90% of the target genes showed age-related changes irrespective of genotype (whose slopes are modified by the teQTLs).

A number of teQTL target genes have important functions in the brain. For example, rs444297 is a teQTL for *EIF4H*, a gene involved in learning and memory in mice (Capossela et al. 2012), as well as Williams syndrome, a disorder whose symptoms can include intellectual disability (Osborne et al. 1996). We found this teQTL in both frontal cortex and cerebellum (Fig. 1D), and its effect was entirely dependent on age, so it was not detectable as an eQTL in either of these brain regions (Supp. Fig. 4). Other teQTL target genes with interesting neurological phenotypes include *AMACR*, a peroxisomal enzyme whose mutation in humans has been observed to result in adult-onset sensory motory neuropathy (Ferdinandusse et al. 2000) and *GAL*, which codes for the neuropeptide galanin, whose immunoreactive fibers hyperinnervate cholinergic neurons in Alzheimer’s disease (Counts et al. 2008).

Interestingly, a number of genes in addition to *EIF4H* were controlled by teQTLs in both the frontal cortex and cerebellum. These two brain regions had significantly more overlap in teQTLs than expected by chance (13 teQTLs in both; permutation p ≤ 1.0×10^-4^), and the effect direction of each allele—either increasing or decreasing expression with age—was also consistent for all 13 (binomial p = 1.2×10^-4^; Supp. Fig. 5A).

After observing this replication across tissues from the same individuals, we tested the teQTLs for replication in an independent data set of 102 dorsolateral prefrontal cortex samples (Colantuoni et al. 2011; referenced here as the “secondary” data set). In this new data set, we again calculated teQTL p-values genome-wide and tested whether there was enrichment for both teQTL significance and matching direction of effect for the teQTLs called in our primary data set (Gibbs et al. 2010). We found significant replication (t-test p = 5.7×10^-5^; 132 pairs have matching direction and p < 0.05; see Methods), despite the primary and secondary data sets differing both in the brain region analyzed and in the expression profiling technology (custom spotted microarrays vs. Illumina BeadChips). We also observed a moderate degree of replication between the primary data set and GTEx frontal cortex data (p = 0.05; see Methods), again despite a number of technical differences (e.g. RNA-seq vs. microarray, and the age distribution of samples).

To further characterize the teQTLs we identified in our primary data set, we investigated the genomic and epigenomic locations of our significant teQTLs in each tissue and the Gene Ontology (GO) categories that were enriched for teQTLs. We found strong enrichment for teQTLs near the TSS of the genes they regulate (p=9.2×10^-13^; Supp. Fig. 6A,B), similar to what has been observed for traditional eQTLs (Supp. Fig. 7-8). Additionally, teQTLs are enriched in non-quiescent regions of the genome when overlapping with brain-specific epigenetic marks (permutation p = 3.5×10^-3^; see Methods; Supp. Fig. 8). Classes of genes enriched for low teQTL p-values included those involved in beta-amyloid binding (p = 7.4×10^-4^) and regulation of cAMP metabolic processes (p = 3.6×10^-4^). The enrichment for beta-amyloid binding is interesting in light of its role in age related disease (Guela et al. 1998). The most enriched GO category was for genes involved in nitrogen compound metabolism (p = 1.6×10^-6^; Supp. Table 3).

### Genetic control of age-related DNA methylation and microRNA expression

One possible molecular mechanism that could explain teQTLs is genetic control of DNA methylation over age, which could in turn affect transcription. To investigate this, we performed a genome-wide scan for temporal methylation QTLs (tmQTLs; Fig. 2A), with the same individuals we used to map teQTLs from the primary data set (Gibbs et al. 2010). At a 20% FDR we identified 14 tmQTL CpG-SNP pairs in the prefrontal cortex, but did not identify any in the cerebellum (Supp. Table 4; 64% of tmQTLs also have mQTL p < 0.05). An example tmQTL in the prefrontal cortex is in the promoter of *USH1C* (encoding harmonin), mutations in which can lead to deafness (Verpy et al. 2000).

**Figure 2.**
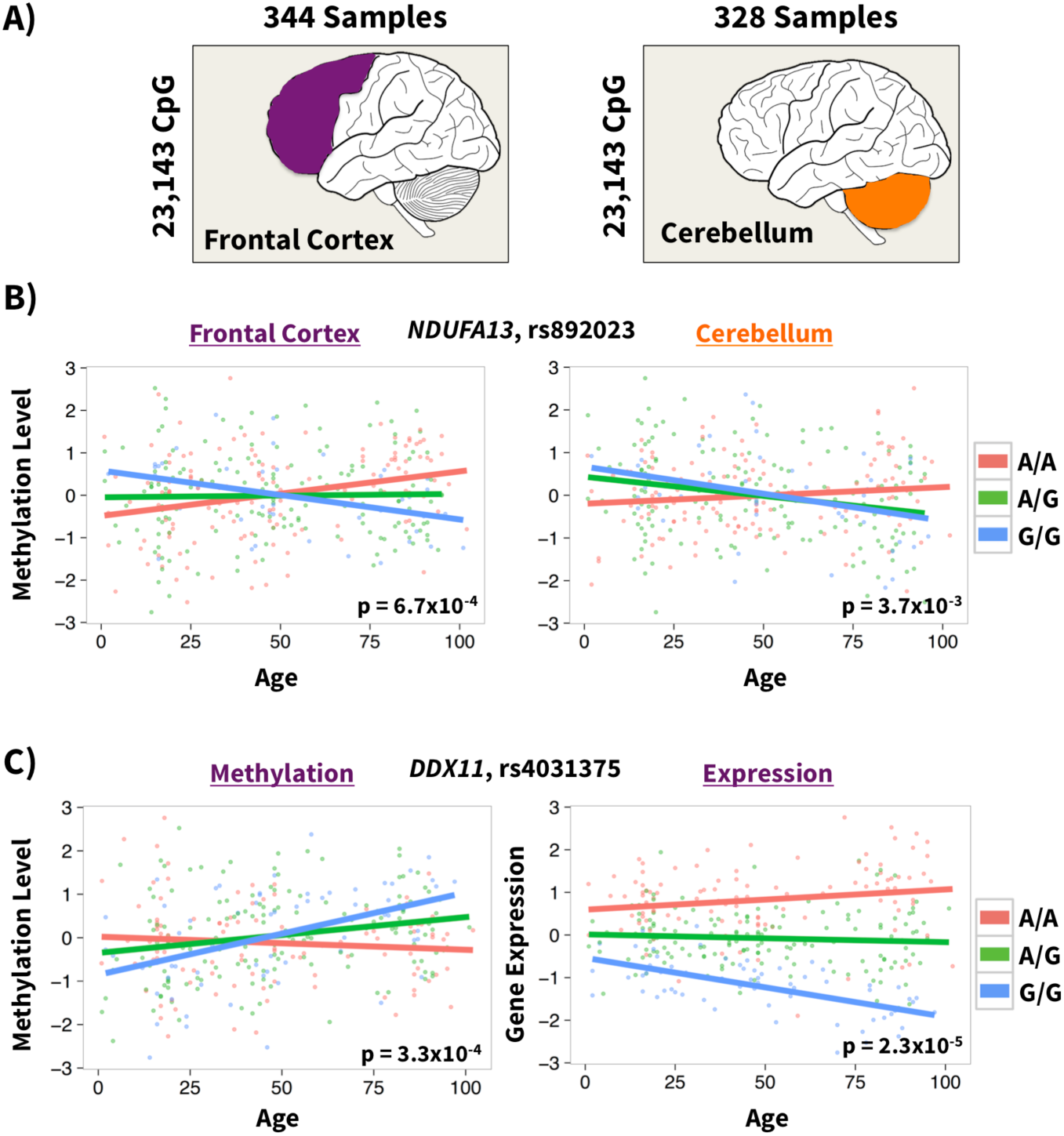
Identification of tmQTLs and overlap with teQTLs. (A) Overview of DNA methylation data. 344 individuals with frontal cortex data (purple) and 328 individuals with cerebellum data (orange) were used in this analysis. (B) Example of a tmQTL present in both tissues. Normalized DNA methylation levels are shown on the y-axis for both the frontal cortex (left; *β* _age*geno_ = -0.01) and the cerebellum (right; *β* _age*geno_ = -0.01) versus age for the gene *NDUFA13*. A linear regression fit for the local SNP rs892023 is also shown as in Fig. 1. (C) Example of a tmQTL that is also a teQTL. Data for the gene *DDX11* is shown as in part (B) for DNA methylation on the left ( *β* _age*geno_ = 0.01) and as in Fig 1 for expression on the right ( *β* _age*geno_ = -0.01) with the local SNP rs4031375.

To validate these tmQTLs, we tested for their replication in the cerebellum samples (see Methods). Even though we did not identify any genome-wide significant tmQTLs in the cerebellum, the frontal cortex tmQTLs showed significantly lower p-values (permutation p ≤ 1×10^-4^) and concordant effect directions (binomial p = 9.54×10^-7^; Supp. Fig. 5B) in cerebellum. Additionally, we observed enrichment of tmQTLs proximal to the CpG sites they regulate (p = 3.7×10^-6^; Supp. Fig. 6C), as has been observed for traditional mQTLs affecting DNA methylation independent of age (Gutierrez-Arcelus et al. 2013). An example tmQTL that replicated across brain regions involves a CpG site in the promoter of *NDUFA13* and the SNP rs892023 (Fig. 2B). *NDUFA13* is part of the mitochondrial respiratory chain complex (CI), whose abundance in neurons has been linked to Parkinson’s disease (Grunewald et al. 2016).

Increased DNA methylation is most often associated with decreased gene expression, though previous mQTL studies have identified almost as many associations in the opposite direction as well (Gutierrez-Arcelus et al. 2013). To explore the relationship of tmQTLs and teQTLs, we tested for directional bias in cases where a teQTL is also a tmQTL. We found a slight enrichment for variants increasing DNA methylation to also decrease mRNA levels (Fisher’s p = 4.5×10^-3^; 64% matching direction, compared to 50% expected by chance), consistent with previous mQTL studies (Gutierrez-Arcelus et al. 2013). An example of a SNP that is both a teQTL and a tmQTL is rs4031375 with the gene *DDX11* (Fig. 2C). *DDX11* is a helicase linked to Warsaw Breakage Syndrome, a disorder whose symptoms include intellectual disability (Sun et al. 2015) and may also affect telomere length (Vasa-Nicotera et al. 2005).

Another potentially important regulator of mRNA levels over age is microRNAs. Using expression data for 561 miRNAs in 116 of the same primary data set individuals, we performed a genome-wide scan for temporal microRNA (tmi) QTLs (Fig. 3A). One miRNA, miR-517*, had a significant tmiQTL in the prefrontal cortex (p = 3.0×10^-4^; Fig. 3B); no miRNAs reached genome-wide significance in the cerebellum. To further characterize the tmiQTL for miR-517* we hypothesized that this SNP may also be a trans-acting teQTL for the mRNA targets of miR-517*, as a downstream consequence of its effect on miR-517*. Among 34 experimentally verified miR-517* targets (Chou et al. 2016) we observed a significant enrichment of trans-teQTLs (p = 1.2×10^-2^). Interestingly, for the majority of these trans-teQTLs, the SNP genotype associated with an increase in miRNA expression over age also increases mRNA expression over age (p = 3.7×10^-3^). The typical model of miRNA action on its targets would be to decrease mRNA expression with increases in miRNA levels, but studies have increasingly shown that when testing across individuals there are many examples of both positive and negative correlations between expression levels of miRNAs and their target mRNAs (Gamazon et al. 2012; Vasudevan et al. 2007). An example of a trans-regulated target of miR-517* is the gene *ALDOA* (Fig. 3C), a glycolytic enzyme implicated as an oncogene in several types of cancer (Ji et al. 2016, Kawai et al. 2017).

**Figure 3.**
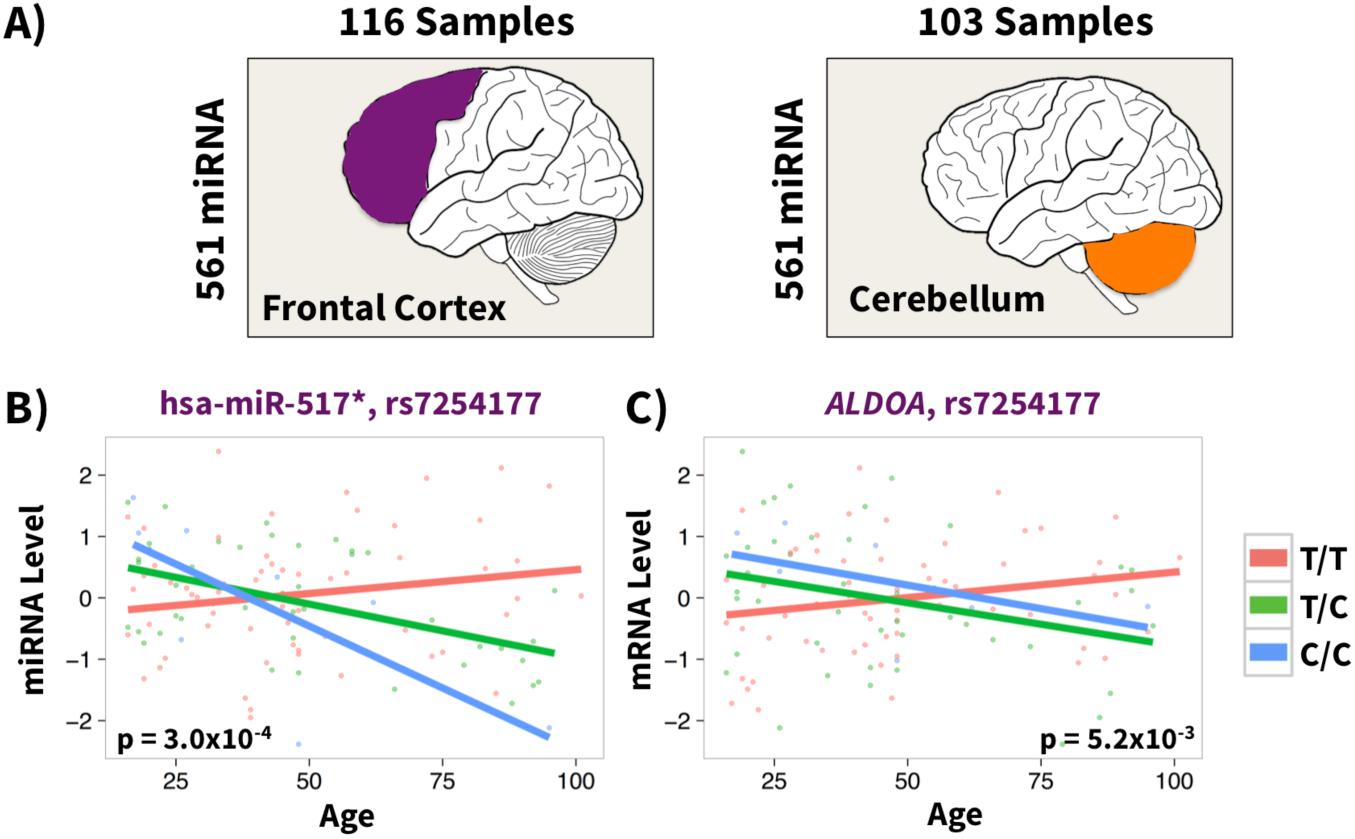
Identification of tmiQTLs in two brain regions. (A) Overview of the miRNA expression data. 116 individuals with frontal cortex data (purple) and 103 individuals with cerebellum data (orange) were used in this analysis. (B) Example of a tmiQTL in the frontal cortex. Normalized miRNA expression level for miR-517* is shown on the y-axis for the frontal cortex with data for the local SNP rs7254177 shown as in Fig. 1 ( *β* _age*geno_ = -0.02). (C) Example of a trans-teQTL for a target of hsa-miR-517*. The gene *ALDOA* is a target of miR-517* and is a trans-teQTL with the local tmiQTL for miR-517* ( *β* _age*geno_ = -0.02). Data plotted as in Fig. 1.

Using a relaxed threshold to evaluate tmiQTLs with consistent effects in both frontal cortex and cerebellum, we found significant replication (p = 2.7×10^-2^; see Methods). One of the tmiQTLs with the strongest replication was miR-9*, the antisense strand of miR-9, a well-studied microRNA that regulates neurogenesis (Coolen et al. 2013).

### The role of teQTLs in complex traits

To assess whether teQTLs are important determinants of complex disease risk, we compared teQTLs discovered in the primary data set with published genome-wide association study (GWAS) results. Specifically, we measured enrichment of trait-associated SNPs in our teQTL SNP set by testing for a shift in the distribution of teQTL p-values for SNPs with a GWAS p-value of 5×10^-8^ or lower (indicating a genome-wide significant trait association). We found a highly significant overall shift in teQTL p-values (t-test p = 2.0×10^-56^), indicating an enrichment for teQTLs among disease-associated loci. To look more broadly at the specific classes of GWAS hits that are enriched among teQTLs, we expanded our analysis to include all nominally significant GWAS hits (p < 0.05 in the GRASP GWAS database; Leslie et al. 2014) and tested for enrichment in each of GRASP’s 5,956 disease categories relative to all disease categories. Although these GWAS hits will include many false positives, previous studies have shown that less stringent cutoffs can greatly increase power to identify genome-wide trends (Richards et al. 2012). We found significant enrichment for 102 categories in the prefrontal cortex teQTLs and 105 in the cerebellar teQTLs. Diseases significant after Bonferroni correction included bipolar disorder, Alzheimer’s disease, Creutzfeldt-Jakob disease, and autism. Additionally, we found that there was significant enrichment more generally for neurological phenotypes (Wilcoxon p = 7.5×10^-3^). The most enriched phenotype category overall in the frontal cortex was height, followed by bipolar disorder (Supp. Tables 5-6).

Many of the phenotype categories identified were also enriched for eQTL associations, so to test the relative strengths of eQTL vs teQTL enrichments we performed the category enrichment test in the same manner as above for eQTLs that we mapped from the same primary data set. Comparing this with the teQTL results, we found that teQTLs were significantly more enriched than eQTLs in neurological traits (permutation p = 1.5×10^-3^; Fig. 4A; see Methods and Supp. Table 7). The neurological trait with the largest over-enrichment for teQTLs was sporadic Creutzfeldt-Jakob disease (teQTL p-value = 2.3×10^-6^; eQTL p-value = 6.4×10^-2^). Other diseases enriched more for teQTLs than for eQTLs included partial epilepsy, autism, and major depressive disorder. Due to linkage disequilibrium (LD), we cannot determine if the same variants are causal for both teQTLs and disease associations; however we do not expect LD to specifically favor teQTLs over eQTLs mapped in the same samples.

**Figure 4.**
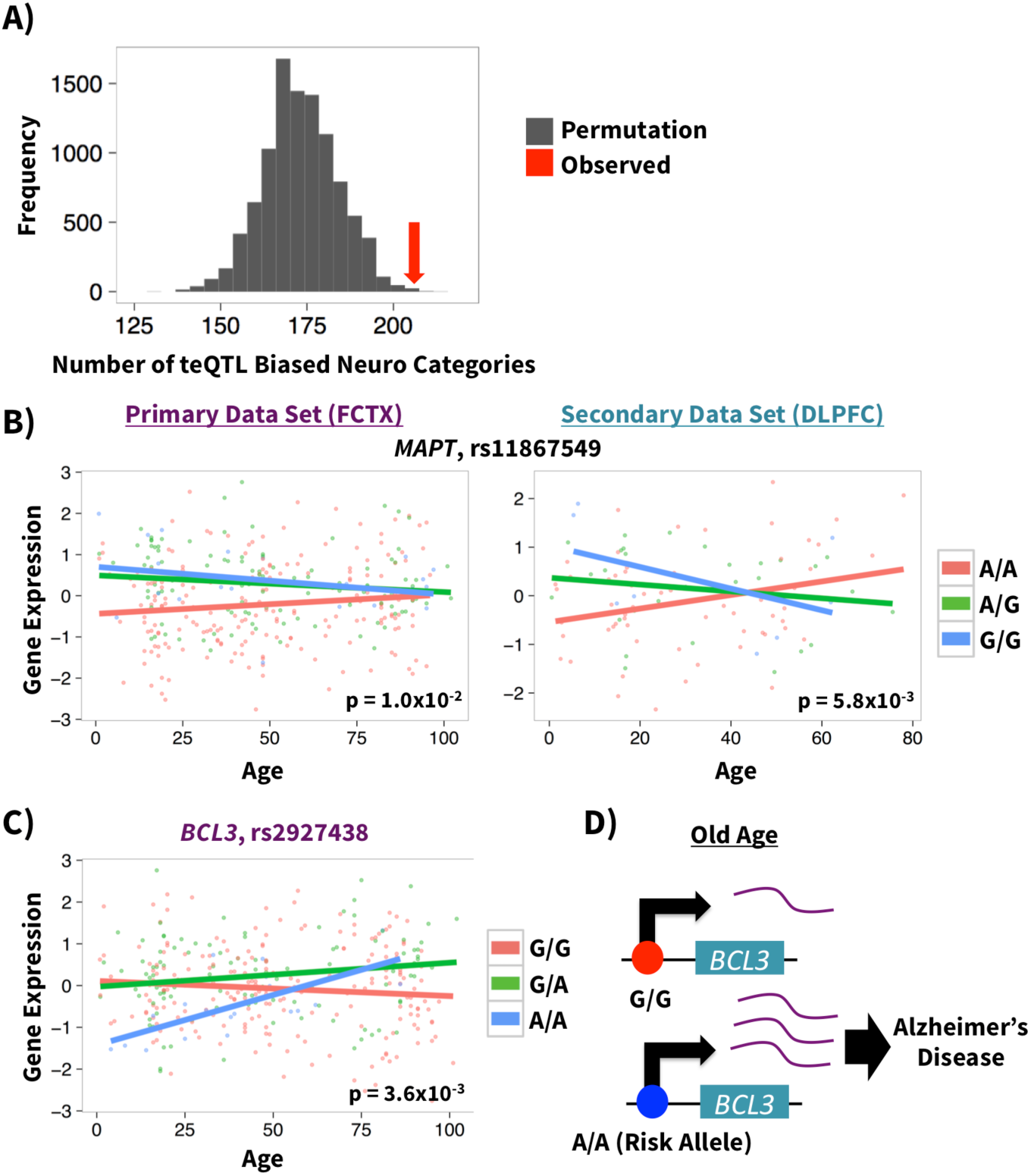
Phenotypic consequences of teQTLs. (A) Neurological categories tend to have stronger teQTL than eQTL enrichments. Permuted biases towards teQTL enrichment in grey histogram, observed bias in red arrow. (B) Example of a teQTL that replicates across data sets and is a GWAS hit for a neurological disorder. The gene *MAPT* is plotted with data for the local SNP rs11867549 as in Fig. 1 (Discovery *β* _age*geno_ = -0.01; Replication *β* _age*geno_ = -0.02). (B) The genome-wide significant Alzheimer’s GWAS SNP rs2927438 is a teQTL for *BCL3*, data plotted as in Fig. 1 (*β* _age*geno_ = 0.01). (C) A proposed mechanism for how the SNP rs2927438 influences Alzheimer’s disease risk in a time-dependent manner.

In other cases, well-established disease-associated variants were also teQTLs. For example, a teQTL for MAPT replicated across the primary and secondary data sets (p ≤ 0.05 in each; risk allele has higher expression early in life) and is a genome-wide significant GWAS hit for progressive supranuclear palsy (PSP; Fig. 4B) (Hoglinger et al. 2011). *MAPT* encodes the microtubule-associated tau proteins, which play a central role in tauopathies including PSP, frontotemporal dementia, and corticobasal degeneration (Baker et al. 1999). Another example of a teQTL (Fig. 4C; p = 3.6×10^-3^) that is also a genome-wide significant GWAS hit is the SNP rs2927438 with the gene *BCL3* (Hollingworth et al. 2011). Late in life the rs2927438 risk allele for Alzheimer’s up-regulates *BCL3* (Fig. 4D), concordant with the *BCL3* up-regulation observed in mouse models of Alzheimer’s disease (Orre et al. 2014).

Other phenotypes with teQTLs include genome-wide significant genetic loci involved in longevity and arthritis. A genome-wide scan for SNPs involved in exceptional longevity identified rs405509, which is also a teQTL with the gene *CLPTM1* (p = 4.8×10^-3^; longevity increasing allele increases expression over age) (Fortney et al. 2015). Mutations in *CLPTM1* can lead to cleft lip and palate in humans (Yoshiura et al. 1998).

## Discussion

In this study we have identified genetic variants controlling three molecular traits—mRNA levels, DNA methylation, and miRNA levels—in an age-dependent manner. We found that this type of genetic control is most pronounced in the human brain, and particularly in the caudate basal ganglia, which is known to play a key role in age-related neurological disease (Stocco et al. 2010). It is also notable that we observed significant sharing of teQTLs across frontal cortex and cerebellum, as well as replication across independent data sets utilizing different cohorts and different expression profiling technologies.

We found that teQTLs are specifically enriched among GWAS variants associated with neurological phenotypes, with stronger enrichment than eQTLs for this class of diseases. These results suggest that teQTLs could play a role in explaining the age-dependency of neurological disease pathogenesis, though further work will be required to test this and to compare their effects with mutations in genes whose age-associated expression changes are not polymorphic. Additionally, some of the non-neurological phenotypes enriched for teQTLs, such as insulin sensitivity and inflammation-related traits, suggest that age-dependent genetic control could play a role in traits associated with aging and longevity more generally (Franceschi and Campisi 2014; Akintola and van Heemst 2015).

The enriched overlap between teQTLs and tmQTLs suggests that teQTLs may sometimes act via DNA methylation, though increases in DNA methylation at a gene’s promoter do not always result in decreased expression as traditional models of gene regulation would suggest (Gutierrez-Arcelus et al. 2013). Similarly, for one tmiQTL we identified evidence for trans-acting regulation of the microRNA’s mRNA targets but not in the direction expected; increasing miR-517* with age was associated with increases in its target mRNAs. Further characterization of this effect will be necessary to determine how this regulation is functioning at the molecular level.

Limitations of the current study include relatively low power to detect teQTLs active very early in life because of our limited sampling of younger ages, a period of rapid development in which teQTLs could be especially likely to affect phenotypes. We also note that while our results may be influenced by changes in cell type abundance over age, this cannot account for our local (likely cis-acting) teQTLs, since any effects caused by cell type abundance would act in trans on genes throughout the genome (see Methods).

Looking ahead, extensions of this analysis could include mapping age-dependent QTLs for chromatin accessibility or other molecular phenotypes, as well as studying the interaction of genotype with other factors. These could include variables intrinsic to each individual, such as body mass index or blood pressure, or extrinsic, such as smoking or sunlight exposure (Knowles et al. 2015; Kita and Fraser 2016). Such studies will be instrumental for our understanding of how the complex interplay between genotype and environment shapes phenotypes across the human lifespan.

## Methods

### Data acquisition and normalization

The primary data set of prefrontal cortex mRNA expression data, CpG methylation, and miRNA expression data for 364, 364, and 143 individuals respectively and cerebellum data for 364, 350, and 143 individuals respectively was downloaded from GEO Accession Number GSE36192 (mRNA), GSE15745 (methylation), GSE15745 (miRNA) and genotype data from dbGaP Study Accession phs000249.v2.p1 (Gibbs et al. 2010; Hernandez et al. 2011). mRNA expression was measured on the Illumina HumanHT-12 V3.0 expression beadchip, CpG methylation data on the Illumina HumanMethylation27 BeadChip, and miRNA expression data on the Illumina Human v1 MicroRNA expression beadchip (Gibbs et al. 2010; Hernandez et al. 2011). Additionally, a secondary data set of dorsolateral prefrontal cortex mRNA expression and genotype data for 269 individuals was downloaded from GEO Accession Number GSE30272 and dbGaP Study Accession phs000417.v1.p1 (Colantuoni et al. 2011). Finally, processed mRNA expression and genotype data on 44 tissues (the GTEx data set; GTEx_Analysis_V6_eQTLInputFiles_geneLevelNormalizedExpressionMatrices.tar.gz) and varying numbers of individuals was downloaded from the GTEx project website (http://www.gtexportal.org/) and dbGaP Study Accession phs00424.v5.p1 (GTEx Consortium 2015). The GTEx sample ages range from 20-70 years old and has data for 19,950 protein coding genes, the primary data set from ages 1-102 years old and has 16,945 genes, and the secondary data set from ages 14 weeks post-conception to 78 years old and has 12,005 genes.

For both primary and secondary data sets, expression measurements were log2 transformed and then all measurements had covariates estimated and removed using the program PEER with default parameters (Stegle et al. 2012) followed by the mapping of the expression and methylation levels to a standard normal distribution for each gene. Additionally for all data sets only samples with Caucasian ethnicity and age greater than or equal to one year, and only autosomal genes, were included in this analysis. We identified 20% of probes in the primary data set and 36% of probes in the secondary data set as having significant (corrected p < 0.05) correlations with age after Bonferroni correction. Additionally, previous studies (Yang et al. 2015) have found genes associated with neurological phenotypes whose expression changes over age in the GTEx data.

The gene and SNP meta data including transcription start sites and SNP locations for the primary and secondary data sets were annotated using Ensembl’s biomaRt package (Durinck et al. 2009) and human genome build GRCh38. GTEx data were annotated using GENCODE release 19 (GRCh37) (Harrow et al. 2012). miRNA locations with hsa-miR IDs were annotated using miRBase (Griffiths-Jones et al. 2006) and those with HS IDs annotated using the genomic locations from Supplementary Table 2 of Berezikov *et al*. (Berezikov et al. 2006). All analyses were performed in R and scripts are available upon request.

### teQTL mapping

Temporal genetic control of expression and methylation was detected by a linear regression model of the form: expression ∼ gender + genotype + age + age x genotype. For the primary and secondary data sets PEER covariates were regressed out beforehand (estimated using PEERset_Nk); for the GTEx analysis the covariates provided by GTEx were included in the interaction model. Permutation-based p-values for each gene were obtained for the interaction term age x genotype through using the software MatrixEQTL to permute the sample labels (Shabalin 2012). To obtain the per-gene p-value all SNPs within 50kb upstream or downstream of the gene TSS, CpG site, or miRNA genomic location (our cis-window) and having MAF > 5% were tested for the interaction term and the minimum p-value across these SNPs was kept as the test statistic. The gene-level empirical p-value is then:

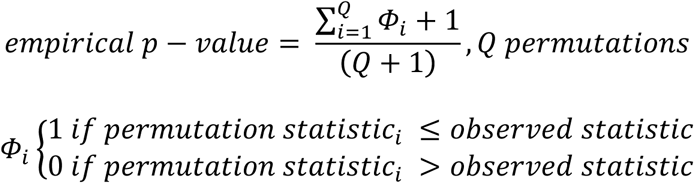

Where 10,000 permutations were performed for all genes, and 10,000 additional permutations were performed for each gene until at least 15 permuted statistics were smaller than the observed statistic, up to a maximum of 600,000 permutations. These gene-level empirical p-values were then converted to local FDR q-values using the q-value R package with default parameters (Storey 2002). We then expanded our list of teQTLs to all SNPs that had a significant association (not only the best SNP) by calculating the empirical p-value for all SNPs in the local window, comparing again to the permuted minimum statistics, resulting in conservative SNP-level p-values. We then mapped these SNP-level empirical p-values to q-values by fitting a loess curve to the gene-level relationship between p-values and q-values and then using this curve to predict the SNP-level q-values from the SNP-level p-values.

As an additional control, we also permuted the gene expression matrices, maintaining the correlation structure between genes, and reran the full teQTL analysis three times using the same pipeline as above. The mean number of teQTLs discovered in each tissue was 0.42 and the median was 0.

SNPs were annotated using the Bioconductor VariantAnnotation package (Obenchain et al. 2014) and SNP epigenome characterization was performed using data from E073 (dorsolateral prefrontal cortex) and E010 (H9 derived neuron culture cells) for the frontal cortex and cerebellum respectively (Roadmap Epigenomics Consortium et al. 2015).

teQTL and tmQTL SNP enrichment near their regulated gene and CpG site respectively was evaluated by a Kolmogorov-Smirnov test for divergence from the uniform distribution. Location outside of quiescent regions was evaluated by comparing the number of observed SNPs outside of this region as defined by Roadmap in E073 cells and comparing this to 1,000 permuted SNPs.

Enrichment for teQTL SNPs in the GTEx brain tissues was calculated by a Wilcoxon test assessing if there is a significant difference in the number of teQTLs (at a 25% FDR) in the brain tissues versus all other tissues. We also treated all brain samples as a single region (taking the mean number of teQTLs across all brain regions), and compared it to 25 other body regions classified as a general tissue type in GTEx. Brain was 11 standard deviations higher than the mean across other regions, suggesting that brain is an outlier even when treated as a single sample. We also redid this calculation using the fixed expression permutation approach described above, which preserves relationships between brain regions, and did not see enrichment for teQTLs in the brain in the randomized data (Wilcoxon p = 0.72).

### Cross-validation and overlap

Results from the primary data set were cross-validated in the secondary data set by testing if the distribution and direction of gene-SNP pair nominal p-values in the secondary data set were significantly shifted for gene-SNP pairs that have a nominal p-value less than 0.05 in the primary data set. Genes were matched across data sets by official gene symbol and SNPs were matched by rs ID. A shift in the p-value distribution and direction was assessed by first checking, for each matched gene-SNP pair, if the sign of the age x genotype interaction coefficient was the same. Then, a one-sided t-test was performed on the following transformed values:

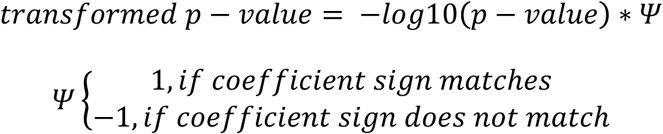

If these transformed p-values are significantly greater than 0 then there is a shift in the secondary data set among the gene-SNP pairs identified in the primary dataset, indicating replication of the primary data set results.

We also validated our results across tissues within the primary data set by counting the number of gene-SNP pairs with −log10 transformed p-values greater than a set threshold (3 for mRNA expression, 1.5 for methylation, 0.4 for miRNA expression) in both the frontal cortex and cerebellum compared to 1,000 random permutations of pairs (matched for both MAF and distance to transcript using four bins for each). Additionally, we performed a binomial test on the number of these pairs that matched direction, where the random expectation is 50% matching.

Enrichment for tmQTLs having the opposite direction of teQTLs was performed by first taking all gene-SNP pairs that had a −log10(p-value) of 2.5 or higher for the tmQTL analysis and using Fisher’s exact test to ask how often these SNPs matched with the same genes had an opposite direction coefficient in the teQTL analysis.

### GWAS enrichment

SNP-level GWAS p-values were downloaded from the GRASP database (Leslie et al. 2014). These GWAS SNPs were then expanded to include SNPs in LD by finding any SNPs with r^2^ > 0.7 and within 500kb of the GWAS SNP based on the 1000 Genomes project EUR population (1000 Genomes Project Consortium et al. 2012). To calculate overall enrichment of teQTLs in GWAS hits, the distribution of primary data set teQTL p-values for SNPs matching a GWAS SNP were compared to the overall distribution of teQTL p-values using a two-sided t-test after regressing out the variance explained by MAF and distance to transcript. To test enrichment for the 5,956 phenotype categories (identified by the GRASP database) within the GWAS SNPs (thresholded to have GWAS p-values < 0.05) first the minimum teQTL p-value for each LD-expanded GWAS SNP was taken. This LD-culled list was then used to compare a specific phenotype category SNP subset to all GWAS SNPs by a t-test for a shift in p-value distribution. These phenotype categories were then Bonferroni corrected and significance assessed. Enrichment for neurological disease associations was evaluated by a Wilcoxon test on the difference in neurological disease teQTL enrichment p-values compared to all categories.

To compare enrichment of teQTLs and eQTLs among neurological disease-associated SNPs, we performed eQTL enrichment calculations in the same manner as above and then classified each neurological category in a binary fashion as either having a stronger teQTL or eQTL enrichment. eQTLs were mapped using the same parameters as teQTLs; eQTL significance was determined by the main effect of genotype on gene expression. 10,000 permutations of the category labels of the enrichment data were then performed and the number of observed enrichments compared to the number of permutation enrichments as described previously to calculate an empirical p-value.

### GO enrichment

Gene ontology enrichment for genes ranked by teQTL p-values was tested in this analysis using the g:Profiler R package with default settings and the “fdr” correction for multiple testing (Reimand et al. 2011).

### Cell type abundance changes are unlikely to explain teQTLs

Any genetic variants affecting relative cell type abundance over age could potentially be teQTLs; however these will always have a *trans-acting* mechanism, affecting genes throughout the genome, rather than just nearby the teQTL variant. In this study we see strong enrichment of teQTL variants near the TSSs of the genes they regulate, strongly suggesting a cis-acting mechanism affecting transcription. Therefore teQTLs driven by changes in cell type composition over age are very unlikely to account for the teQTLs we have detected.

### miRNA target analysis

For 116 (frontal cortex) and 103 (cerebellum) individuals analyzed for teQTLs we also had miRNA expression data available from the primary data set (GSE15745). miRNA targets were downloaded from miRTarBase (Chou et al. 2016). Significant local tmiQTLs were then checked for having enrichment for trans-teQTLs (calculated using the same pipeline as cis-teQTLs) with the targets of the relevant miRNA by a two-sided Wilcoxon test (background of random expressed genes) and also for biased direction of the effect among the targets using a two-sided binomial test (background is percent of expressed genes with matched direction).

## Acknowledgements

We thank members of the Fraser lab for helpful feedback and advice. This work was supported by NIH grant 2R01GM097171-05A1. TM is supported by an NSF predoctoral fellowship.

## Table Legends

**Table S1** List of teQTLs significant at 20% FDR in the frontal cortex

**Table S2** List of teQTLs significant at 20% FDR in the cerebellum

**Table S3** List of top 40 gene ontology results in the frontal cortex

**Table S4** List of tmQTLs significant at 20% FDR in the frontal cortex

**Table S5** List of GWAS categories with significant teQTL p-value shift in the frontal cortex

**Table S6** List of GWAS categories with significant teQTL p-value shift in the cerebellum

**Table S7** List of top 40 GWAS categories with most teQTL over eQTL enrichment and top 40 GWAS categories with most eQTL over teQTL enrichment

